# A neural marker of eye contact highly impaired in autism spectrum disorder

**DOI:** 10.1101/2021.03.29.433074

**Authors:** Guillaume Lio, Martina Corazzol, Roberta Fadda, Giuseppe Doneddu, Caroline Demily, Angela Sirigu

## Abstract

Attention to faces and eye contact are key behaviors for establishing social bonds in humans. In Autism Spectrum Disorders (ASD) a neurodevelopmental disturbance characterized by poor communication skills, impaired face processing and gaze avoidance are critical clinical features for its diagnosis. The biological alterations underlying these impairments are not clear yet. Using high-density electroencephalography coupled with multi-variate pattern classification and group blind source separation methods we searched for face- and face components-related neural signals that could best discriminate neurotypical and ASD visual processing. First, we isolated a face-specific neural signal in the superior temporal sulcus peaking at 240ms after stimulus onset. A machine learning algorithm applied on the extracted neural component reached 74% decoding accuracy at the same latencies, dissociating the neurotypical population from ASD subjects in whom this signal was weak. Further, by manipulating attention to face parts we found that the signal-evoked power in neurotypical subjects varied as a function of the distance of the eyes in the face stimulus with respect to the viewers’ fovea, i.e. it was strongest when the eyes were projected on the fovea and weakest when projected in the retinal periphery. Such selective face and face-components neural modulations were not found in ASD individuals although they showed typical early face related P100 and the N170 signals. These findings show that dedicated cortical mechanisms related to face perception set neural priority for attention to eyes and that these mechanisms are altered in individuals with ASD.

## Introduction

Autism spectrum disorder (ASD) is a lifelong neuro-developmental disorder that begins in early childhood. Despite the fact that the first signs of autistic behaviors are observable during the first months of life^1,2^ in current medical practice ASD is diagnosed far later, around the age of three^3^. Although numerous biological and environmental factors have been considered in the study of its pathology^4^, a combination of behavioral observations and clinical interviews remains the principal method of diagnosis. These assess the heterogeneity of ASD deficits which impact social communication, social reciprocity and include repetitive and stereotyped behaviors^5^. Indeed, recent epidemiologic studies show that up to 1 in every 54 individuals are affected by ASD, but estimates can differ greatly between countries^6^.

Despite the absence of diagnostic advances, numerous cortical abnormalities linked with autistic syndrome have been detected over the past decade. Approaches have ranged from post-mortem studies^7–10^ to the use of imaging techniques such as resting functional Positron Emission Tomography (PET)^11,12^, Single-Photon Emission Computed Tomography (SPECT)^13^ and structural MRI (e.g.^14^ for a review). Although these neuroimaging studies have made substantial progress, the discovery of a reliable and clinically useful biomarker of autism remains elusive^4,15,16^. Despite evident advantages, the use of electro-encephalographic (EEG) techniques has been somewhat underexploited as a tool in achieving this goal.

In contrast to some neuroimaging approach, high density EEG is a noninvasive technique which makes it particularly convenient for longitudinal studies of younger children and adolescents. In addition, advances in signal processing means that this method is particularly amenable to single subject and single trial analyses enabling a more nuanced characterization of critical electrophysiological processes (see e.g.^17–19^). Moreover, cortical and synaptic abnormalities commonly found in ASD patients^7–10,20^ are highly receptive to being correlated with functional modification of post-synaptic activities. These are, therefore, phenomena that can be captured on the scalp using EEG techniques. As a proof-of concept, Bosl and colleagues^21^ demonstrated that the risk status for autism can be reasonably well predicted (≈80% accuracy) by spontaneous EEG (see also^15,22,23^ for discussions on the relevance of such biomarkers for good clinical practice).

In the field of EEG, we have witnessed the growth of an extensive literature on electroencephalographic markers of face processing^24–26^. In addition we have learned the critical role that gaze plays in social interactions^27,28^ and how ASD patients show early impairment in spontaneous eye contact behaviors^1,2,29–31^. That is why many studies have focused on scalp potentials evoked by face stimuli in the hope of detecting and isolating some critical neuro-physiological correlates that might signpost the emergence of autistic phenotypes. Yet, efforts to reveal difference between patients and neurotypical controls amongst the cascade of events that contribute to visually evoked potentials (VEP) have, so far, proven inconclusive. Meta-analyses present conflicting results^32,33^ and in a recent highly controlled study Apicella and colleagues^34^ failed to find any significant differences in early face-sensitive ERPs between neurotypical controls and children with high functioning ASD.

We have identified two key issues that have confounded previous attempts in detecting ASD selective activity in scalp ERPs. The first issue relates to the difficulty involved in extracting meaningful information from an overall signal recorded in the scalp. This is due the fact that such a signal is composed of a mixture of electric fields broadcasted from many cortical sources with highly correlated activity^26,35,36^. The second issue arises from the kind of stimuli generally used in neuroimaging studies. Since low level visual features in face stimuli can have a significant effect on the ERPs observed on the scalp^37–39^ it has been necessary to adapt previous images to be highly controlled and stereotyped. Such experimental prerequisites, however, also have the adverse effect of compromising the natural potential of faces to simulate social interaction. This calls for future studies to employ more ecologically valid and naturalistic social stimuli, so that the complex profile of ASD can be more fully captured^40,41^.

In the present study, taking these two major issues, we investigated the electrophysiological correlates of neurotypical face and -face components- processing. Moreover we explored how these processes are altered within the autism syndrome^2^.

To this aim, we first designed a simple eye-contact task in which controlled face stimuli with natural characteristics preserved were presented at a short inter-personal distance to both ASD individuals and a healthy control population. Then, for offline signal processing, we selected advanced linear decomposition techniques^42^, namely multi-variate pattern classification (MVPC)^43^ and group blind source separation (gBSS)^44,45^. These methods are particularly suitable to isolate among the evoked activities those that best distinguish the studied populations. By doing so, we were able to detect a specific pattern of evoked activities approximately 240ms after stimulus onset, in the most anterior regions of the ventral visual stream. Crucially, we found that these activities were critically impaired in the ASD population. This result suggests atypically high-level perceptual processing of face stimuli in superior temporal regions for ASD individuals compared to controls participants. Next, using the spatial pattern from the previously identified source we extended our initial approach to two replication studies, performed in new groups of adults, young neurotypical subjects and ASD subjects, which we could monitor the single trial dynamic of the superior temporal cortex evoked activity. This novel strategy allowed us to map how the recovered electrophysiological source was modulated in relation to the particular face region focused on by the subject. By using such procedure, we were able to highlights the critical role of the eye region in the emergence of this evoked activity. Our findings and the methods described, offer up new avenues to non-invasively investigate the neurodevelopment of social perception and its abnormalities across a range of neuropsychiatric conditions.

## Results

### Experiment 1

Previous work has shown that, in the context of eye contact, individuals with ASD process face stimuli differently to neurotypical controls^29,46^. The purpose of our first experiment was to establish the temporal point at which electrophysiological differences in the way that ASD patients and neurotypical control subjects process face stimuli are maximal. To achieve this initial aim, we designed a simple face presentation task and used standard topographical analysis and multivariate classification techniques. In the task, we presented natural faces with neutral expressions while neurotypical subjects (N= 10) and ASD participants (N= 13) were instructed to focus on a fixation cross located at a point on the screen corresponding to the space situated between the two eyes of each face stimuli (Fig 1). The size and the distance of face pictures were controlled in such a way to simulate an eye-contact between the viewer and each image. Specifically, interpersonal distance was controlled in reference to two by two interpersonal space that is classically observed between two individuals in a naturalistic setting. Here, we simulated a social interaction at a relatively closer distance of ~56 cm^47,48^.

**Figure. 1.**
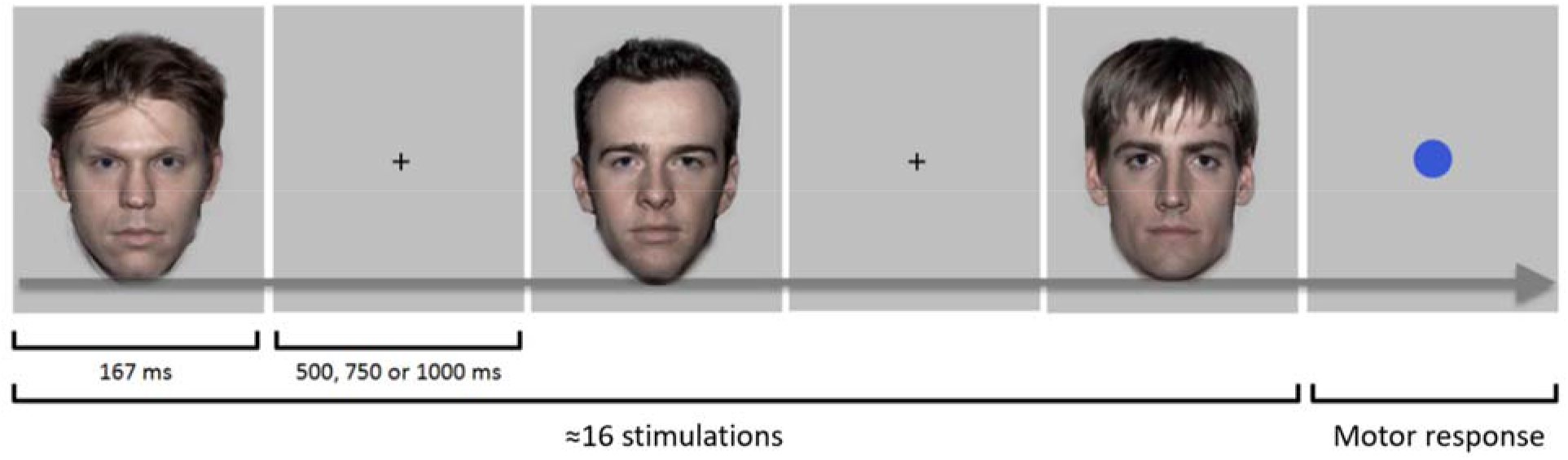
Stimuli presentation experiment 1. Simple face task. Natural faces with neutral expressions were presented for 167ms while subjects were instructed to focus on a fixation cross. In order to simulate a direct-gaze behavior at a relatively close interpersonal distance, face stimuli were large in size and the fixation cross was located on the screen at the same level as the eyes. To maintain the attention of subjects, a blue dot was randomly presented instead of a face stimulus. Subjects were instructed to press a button when the dot appeared.

### Preliminary electrodes space analysis

Both ASD participants and neurotypical controls showed the typical early surface ERPs namely the P100 and the N170 components (Fig 2B; Fig S1). These are classically observed in face evoked ERP studies^24^. No significant differences between the two populations were revealed for these two components by either the cluster permutation test (p>0.05) or by separated Wilcoxon rank sum tests (p>0.05, uncorrected for multiple comparisons). This preliminary result was not surprising in light of previously reported observations in a meta-analysis^32,33^ and a series of highly controlled studies^34^ which suggest that early differences in face evoked potentials are highly dependent on the absence of a fixation cross during the task.

**Figure. 2.**
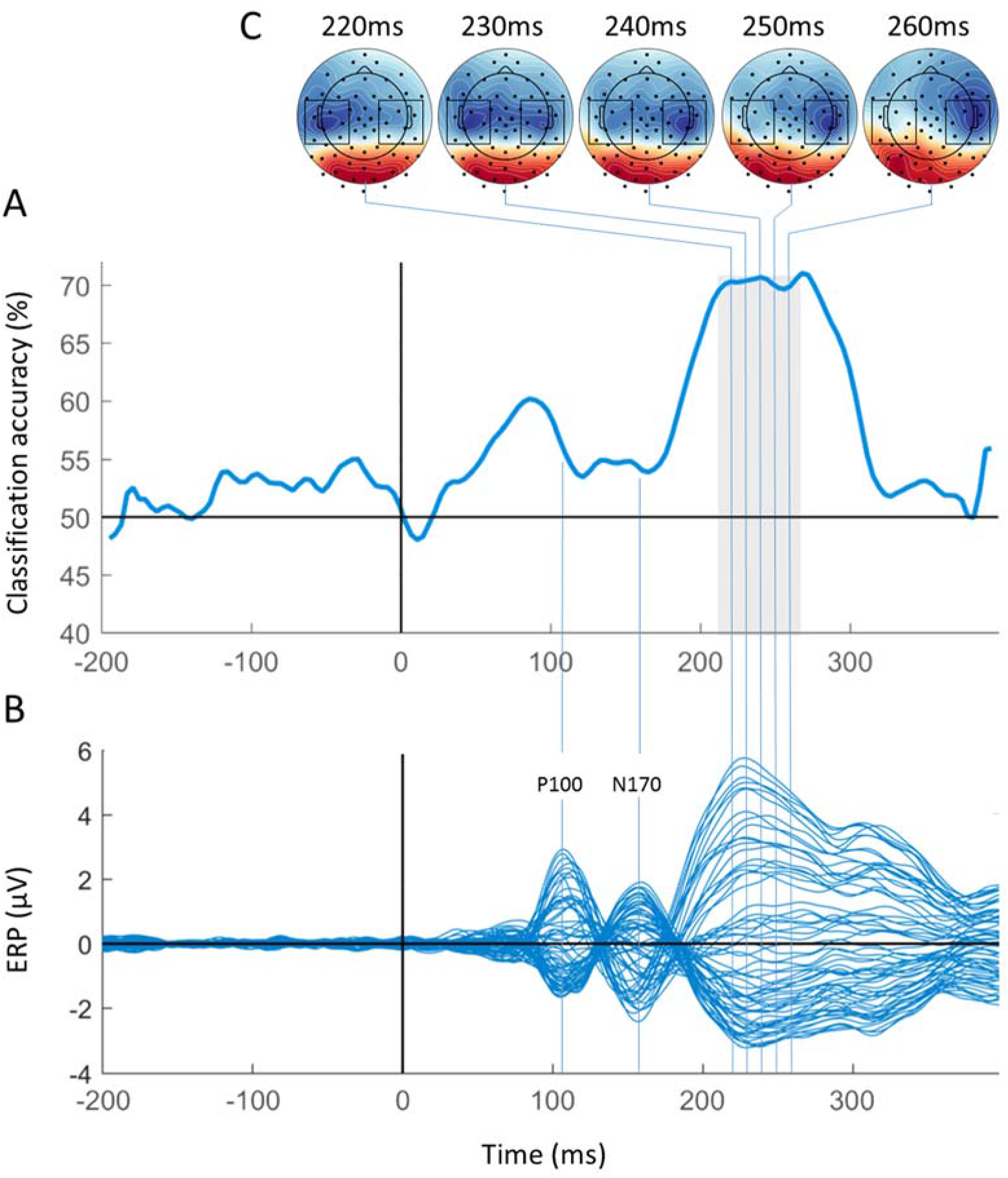
Multivariate pattern decoding of the ASD phenotype. **A**- Time course of the accuracy of the classification of ASD patients and control subjects. The grey area represents the significant time-cluster where differences in the evoked power can be assessed ([220ms 260ms] cluster - p<0.05 cluster corrected). The classification accuracy reached a maximum during this time period (≈70%). **B**- Butterfly plot of the grand average evoked activity recorded on the scalp for the 128 electrodes. The classical P100 and N170 components are well visible but no significant differences were detected. Significant activities were detected later, when multiple parallel processes were triggered for processing the face stimulus. **C**- Spatial patterns that provide the best classification results, estimated by the minimum noise forward model method. Recovered topographies suggest an involvement of multiple sources along the ventral visual pathway. Black squares emphasize the local minima off the patterns, located on the middle upper part of temporal regions.

Both topographical analysis and multivariate pattern classification did reveal differences in evoked scalp potentials during a later time-period. Specifically, topographical analysis yielded significant cluster between 220ms and 260ms post-stimulus onset (CTRL/ASD contrast, time cluster [220ms 260ms], p<0.05 cluster corrected – Fig. S2) while multivariate pattern classification produced similar results. For this technique a bootstrap estimate of subjects’ classification accuracy was calculated for each time sample of the analyzed epochs. To this aim, we trained multiple linear support vector machines (SVM) with leave one-out cross validation to identify the ASD subjects among the “general population”. This procedure yields a time-course of decoding accuracy that can be used to temporally localize the onset, then the maximal observable dissociation in the electrophysiological processing of face stimuli between the two classes of subjects (Fig 2A). Prior to stimulus presentation and up to 200ms after, the grand average decoding accuracy was not significantly different than chance level (50% accuracy). However, classification results rose sharply in the time period between 220ms and 260ms to reach an average level of 70% of accuracy (non-parametric permutation test p<0.05) (Fig. 2A 2B). The spatial patterns related to these results in this time period were then estimated by the minimum noise forward model method^42^ to produce complex topographies (Fig. 2C) consistent with distributed evoked activities along the visual ventral stream. It should be noted that face-evoked activities observed at the scalp surface come about through the mixing of multiple correlated sources indexing many parallel processes^35,36^. As a consequence, we must interpret the location of phenomena identified on scalp potentials with cautions^18,49,50^. In order to address this issue, recent advances in EEG signal processing have been used to decompose the observed scalp signal in independent components (ICs).

### Blind sources space analysis

We implemented Group Blind Source Separation (gBSS) using the Second Order Blind Identification (SOBI) algorithm as opposed to other algorithms due to its robustness with respect to inter-individual/inter-trial variability^44,45^ and its ability to recover highly correlated neural sources^51,52^. This last point was particularly crucial given that early evoked activities in different cortical locations are highly correlated^35,36^.

Following Group Blind Source separation, we observed multiples components in the dorsal and in the ventral visual pathways. Among them, a cluster-permutation test at the source level revealed only one significant source (Fig 3) (Independent Component 15 - CTRL/ASD contrast, time cluster [200ms 300ms], p<0.05 cluster corrected). The identified Independent Component (IC) represented a large evoked activity for neurotypical subjects, which was maximal approximately ~240ms after the stimulus onset. This same pattern, however, could not be consistently found in the ASD population (Fig 3B). The topography of the source consisted of a well-defined configuration at two dipoles suggesting a cortical localization in the temporal gyri and/or sulci (Fig 3C). sLORETA distributed sources localization of the IC showed maximal activity around the lateral fissure (MNI coordinates = X: +65 −65 Y: −20 Z: 10) and a local maximal activity around the inferior temporal sulcus (MNI coordinates = X:+60 −60 Y: −40 Z: −20) (Fig 3D). Taken together the sLORETA current source density in combination with the directions of the two dipole scalp topography and the timing of the evoked activity (240 ms after the stimulus onset) suggest a probable localization of the IC in superior temporal regions as opposed to other more posterior face specific sites (e.g. fusiform gyrus and occipito-lateral regions) which are characterized by earlier peak latencies^25,36^. Finally, we wanted to compare the time course of classification accuracy derived at the source level (Fig 3A) with the time course of classification accuracy estimated in the electrodes space as produced by evoked spatial patterns (Fig 2A, 3A). To achieve this, we performed a second multivariate pattern analysis. This procedure was identical to the preliminary analysis, except that only evoked power at the source level was used as an input to the classification algorithm. In terms of linear separability, we observed differences in evoked power between ASD subjects and the control population in the superior temporal regions at ~240ms after the stimulus onset, reaching a peak of 74% of classification accuracy (non-parametric permutation test p<0.05).

**Figure. 3.**
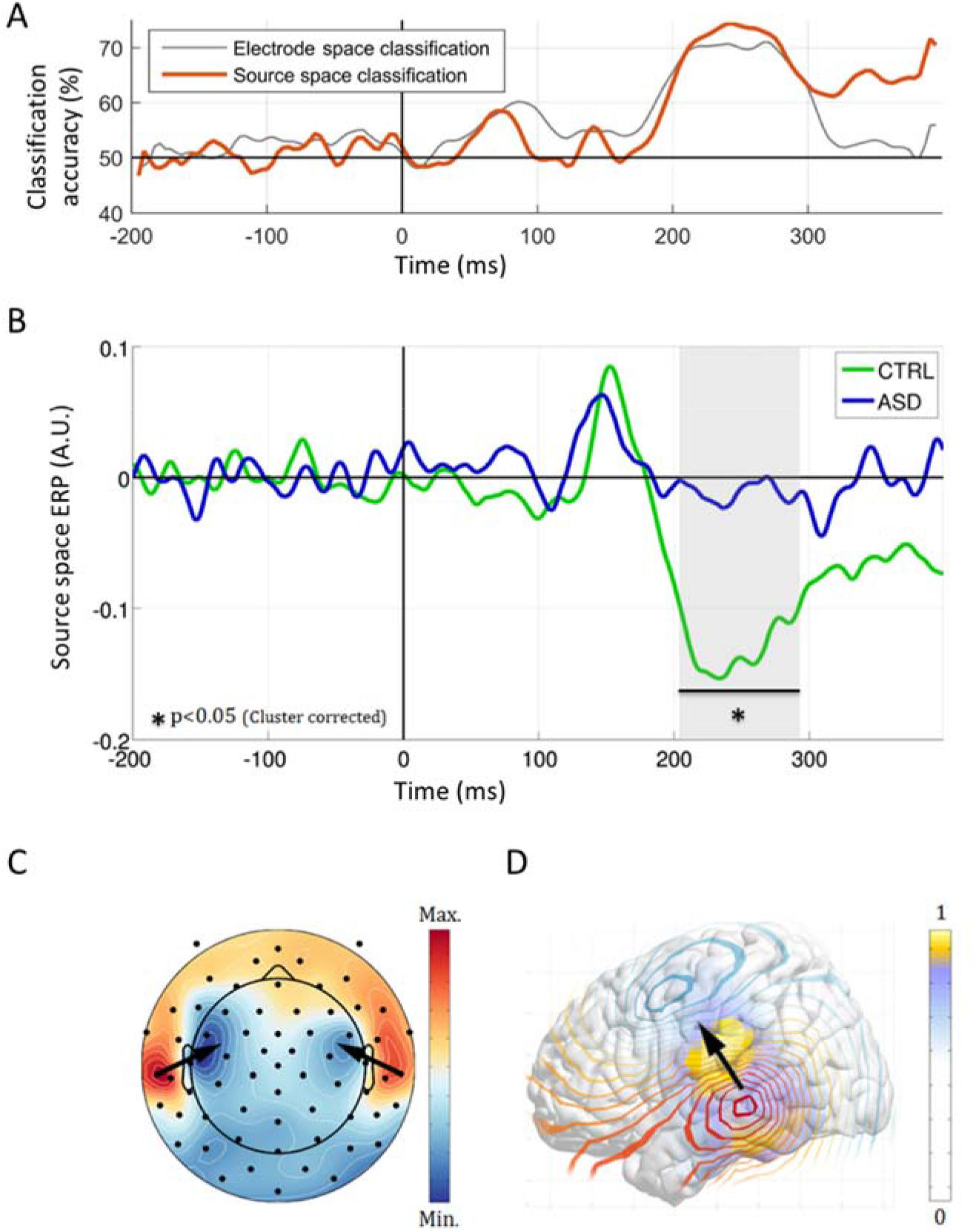
Impaired superior temporal activity in Autism Spectrum Disorder. Autism Spectrum Disorder (ASD) impaired evoked activity during neutral faces presentation, detected after group blind source separation (Independent Component 15). **A**- Time-course of accuracy of the classification of ASD patients and control subjects at the source level, compared to the time-course of classification accuracy estimated in the electrode space. Classification accuracy reaches a peak of 74%, ~240ms after the presentation of the face stimulus. **B**- Evoked group activity at the source level after group blind source separation. Control subjects show a prominent evoked activity between 200 and 300ms after the stimulus onset that is not observable in ASD patients (p<0.05 corrected. Maximum ≈240ms after the stimulus onset). **C**- Scalp topography of the detected independent component (IC). The considered source presents a characteristic bilateral topography. **D**- sLORETA distributed sources localization of the IC. This estimated tomography, located bilaterally on the middle part of the temporal cortex, had a maximum around the lateral fissure and a local maximum around the inferior temporal sulcus. Combined with the directions of the two dipoles on the scalp topography and the latency of the evoked activity in the region, this information strongly suggests a location of the component in superior temporal regions.

In this first experiment, we employed ecologically valid stimuli, thus preserving the social content and implemented signal processing procedure designed to dissociate inter-mixed evoked activities. Using this method, we have pinpoint a neural function which shows a specific impairment among the ASD population. We make this interpretation in light of the established association of the bilateral superior temporal regions in social cognition as well as their implication in neural dysfunction underlying ASD^53,54^. Consistent with this interpretation we postulate that the neural process observed in the control population is strongly associated with the eye contact component simulated in the current task. This generates the logical next question of whether this evoked activity remains if subject attention is directed toward a different face area? In other words, does the simulation of an ‘autistic gaze aversion behavior’ reduce the readable social content projected on to the fovea and, as a consequence, the evoked activities in superior temporal regions?

To address these questions, we designed a replication study which aimed to extract from the previously identified cortical region the single trial dynamic of the activity evoked by face stimuli.

### Experiment 2

In this second study, we configured the experimental setup that we could assess single trial variation of the previously identified component as a function of the face part focused on by neurotypical subjects. Fourteen naïve subjects who had not took part in the first experiment participated in the protocol. We deemed this new paradigm to be essential for two reasons: First, if we find that the observed neural response is modulated by the face part that is being focused on, especially the eye region, then we can be hypothesized that the previously identified neural source is highly related to the social significance of the presented stimulus. In other words, the targeted activity would be correlated to the subjective quantity of social information that can be extracted by the subject from the region presented on to the fovea. Second, if the identified neural source presents a characteristic and reproducible modulation that can be measured at the single-subject level, then this could be a potentially beneficial target for future use in longitudinal studies on toddlers. Moreover, it could also be useful in the development of biomarkers of the social deficits that are characteristic of the autism spectrum disorder.

To this end, we controlled in proportion, position and luminance distribution of face stimuli used in the first experiment. This enabled us to assess and record the exact location on the face that was focused on by the subject for each trial (Fig. 4, Fig. S1). We presented a total of 2800 stimuli to each participant so that each part of each face stimulus was repeatedly observed. Then, we applied spatial filtering using a minimum variance beamformer (SAM^55^) on single-trial data in order to quantify the evoked activity in the previously identified temporal regions. Finally, we combined Gaussian kernel density estimations and non-parametric permutation tests to precisely reveal regions in the face that evoke the most prominent activities during the previously identified post-stimulus time period [200ms, 300ms].

**Figure 4.**
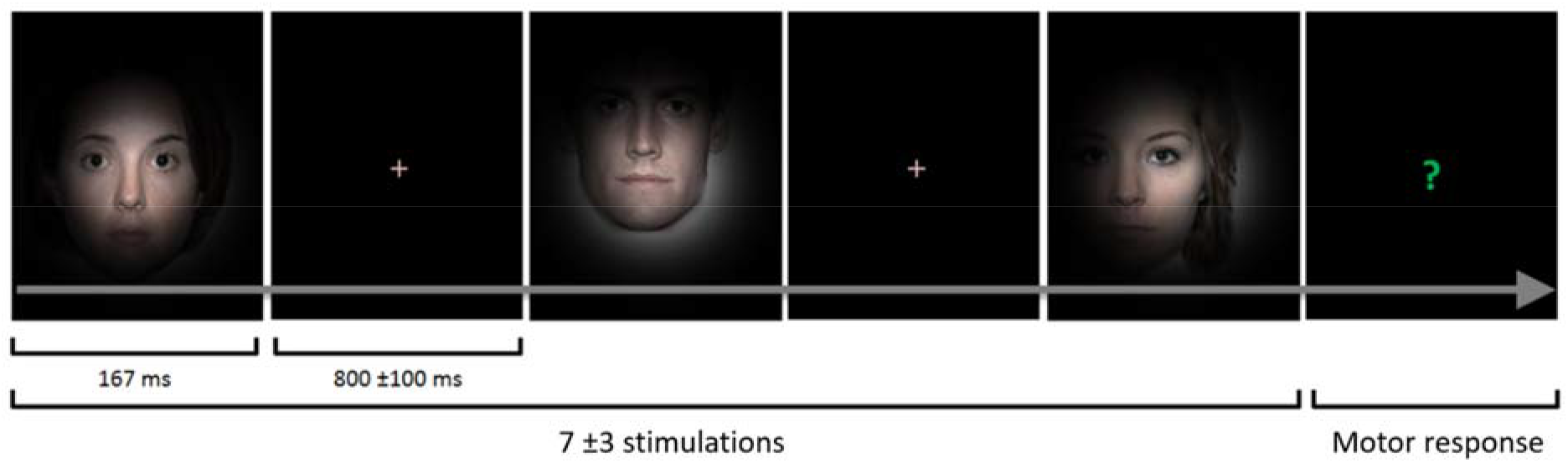
Stimuli presentation experiment 2. Face presentation with a focus on a randomly determined face location. Subjects were instructed to focus on a fixation cross, then a face stimulus multiplied by a Gaussian apodization windows was presented at the center of the screen (FWHM = 10°). The face area to be focused on was randomly drawn with a uniform distribution while the apodization windows is always centered on the fixation cross area. Thus, each stimulus had approximatively the same luminance distribution. When a), a question mark appeared randomly (every 7±3 trials, the subject was instructed to determine the gender of the face stimulus from the preceding trial (left/right button press).

At the single-subject level, different patterns of activation were observed but we found that for the majority of subjects, the global maxima were consistently located on the eye region (11/14 subjects – ≈78% - permutation test, p<0.05) (Fig S4). These preliminary results were found also at the group-level and further confirmed that the neural activity was related to subjects’ attention around the eyes region. Indeed, Gaussian kernel density estimation for the whole population produced a characteristic pattern composed of significant activity encompassing both eyes of the face stimuli, with a maximum activity when subjects were focusing their attention right between the two eyes (permutation test, p<0.05 FWER corrected) (Fig 5A, 5B). Notably, activity in the cortical sensitivity map highlights the same area of the face that neurotypical subjects naturally focus on during social interactions and that individuals with ASD have generally been found to avoid^32,56^ from the first months of life^2^. In other words, the modulation of the evoked power of the bilateral-temporal source identified in the first experiment is highly correlated to eye-contact behavior.

**Figure 5.**
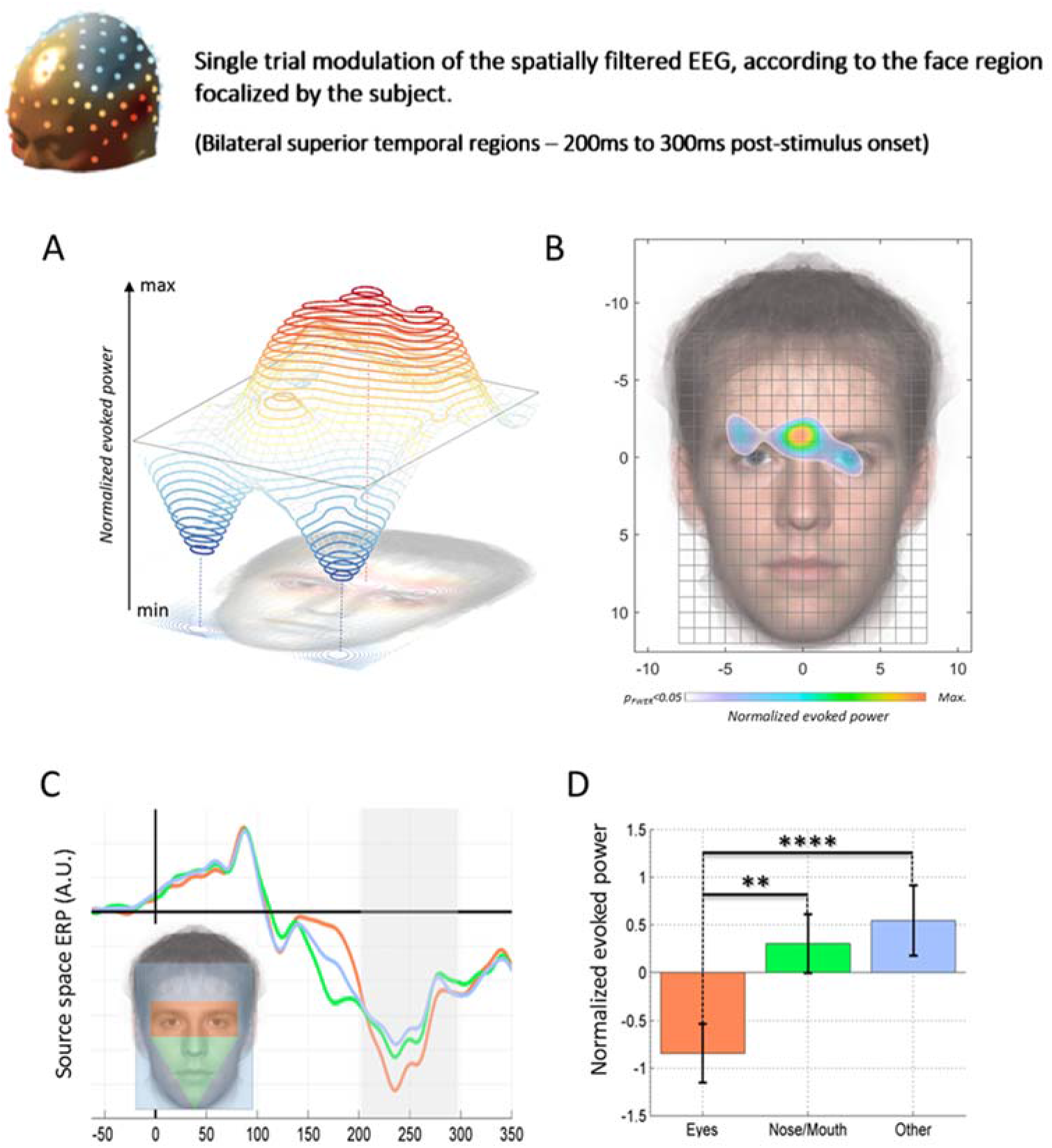
Evoked activity in superior temporal regions is eye-sensitive. Single trial modulation of the spatially filtered EEG, according to the face region focused on by subjects. For each subject the evoked activity in superior temporal regions, 200ms to 300ms post-stimulus, is normalized and mapped as a probability density function of cortical activation. **A**- Group average of the Normalized Gaussian Kernel Density mapping of the evoked activity in the superior temporal regions. Evoked power was maximal when subject’s foveal vision was located in the Eyes/Eyebrows region while two local minima can be observed in the lower part of the tested area, when the line of sight is pointing outside of the face picture. **B**- Statistical non-parametric mapping of the cortical evoked activity (permutation test, p<0.05 FWER corrected). Only the region around the eyes remained significant, with a maximum sensitivity when subjects were focusing in a particular region, between the two eyes. Notably, such a cortical sensitivity map highlights the face parts that neurotypical subjects naturally tend to focus on during a social interactions and that autism patients tend to avoid. (Pelphrey et al. 2002, Simmons et al. 2009). **C**- Regions of interest (ROI) analysis: Three ROIs were selected for analysis. (1) A rectangular area covering the region around the eyes, (2) a triangular area encompassing the nose and the mouth and (3) a third area including the remaining tested areas. These regions were the ones most gazed on by neurotypical subjects in accordance with stereotyped gaze strategies observed during social interactions s by previous studies (Pelphrey et al. 2002). The three waveforms represent the group-averaged evoked activity in the superior temporal regions, for the three ROIs considered. The recovered source reflects the same negativity of the source recovered in the first experiment, in the 200ms – 300ms time period, reaching a maximum approximately 240ms after the stimulus onset. **D**- The eye region showed a marked higher activity relative to the nose/mouth area (p<0.01, FWER corrected) and relative to the other face area (p<0.0001, FWER corrected). No significant differences were found between the nose/mouth area and the ‘other / not internal face features’ area (p>0.05, FWER corrected).

Nonetheless, it is possible that our application of non-parametric permutation tests corrected for multiple comparison could be overly conservative. Other areas of the face that could not be captured by our initial approach could also present significant modulations of the evoked neural response of superior temporal regions. These other areas can be important for social interaction, perception of facial expression and can also contain information about ‘implied biological motion’^53,57,58^ (e.g. the mouth). Therefore, a second analysis using three facial regions of interest (ROI) (Fig 5C) encompassing the eyes region, the lower face features and the rest of the face was also carried out. We observed that grand average evoked power for these three ROIs consisted of a pronounced negative activity occurring from 200ms to 300ms post-stimulus, with a peak-latency at 240ms. Such a pattern showed a remarkable consistency with the electrophysiological process recovered in the first experiment for the neurotypical population (Fig 5C). A Kruskall-Wallis non-parametric test revealed on a global effect of the face ROIs on this signal (χ^2^(2, *N*=14)=20, p<0.0001) and pair-wise post-hoc comparisons confirmed the singularity of the eye region for the observed component: That is evoked activity (200ms to 300ms after the stimulus onset) was significantly higher when subjects focused on the eyes region than when they focused on the mouth/nose area (p<0.01, FWER corrected) and other face regions (p<0.0001, FWER corrected). We found no significant differences in evoked activity between the mouth/nose region and the non-face features area.

In sum, these results suggest that the presentation of naturalistic social stimuli in combination with recent advances in scalp electrophysiological signal processing form a powerful tool in non-invasively and precisely pinpointing in time and in space a critical eye-sensitive electrophysiological process that is particularly impaired in individuals with ASD.

### Experiment 3

Next, in Experiment 3 we implemented a task which was identical to that from the Experiment 2. However, in this third study, we reduced the number of trials by half so that it could be comfortably performed by children (N= 14, mean age 11.6yrs) and a new group of young adults with ASD (N=14, mean age 20yrs). As a tradeoff, however, the spatial resolution generated by the sensitivity maps of the STS evoked responses had to be reduced to obtain a sufficient number of trials per region of the face tested. The results of this third study are presented in Figure 6. We found that both sets of neurotypical populations produced the same response pattern. Specifically, neurotypical subjects showed significant evoked activity in the eye and eyebrow area (p<0.05 FWER corrected) that gradually decreased to reach two local minima on the lower left and right parts, where the area focus is outside the face. In the ASD population, at the group level, we recovered a particular response pattern from this adapted design whereby the maximum and significant activity (p<0.05 FWER corrected) occurred in the middle of the face over the nose region. We also observed the same pattern of decreased activity when the center of the region focused on was outside the face. These results confirm the modulation of activities evoked in the STS with a maximum occurring for neurotypical populations, when the region observed is at the eye and eyebrow level. Since these regions are particularly rich in information essential to the proper functioning of social cognition abilities, an immediate interpretation would be that this observed activity reflects the propensity of subjects to perform a social analysis of presented faces. Interestingly, we also found modulated – albeit atypical - activity in our second population of individuals with autism spectrum disorders. Although not generalizable to the overall patients’ population we observed that, within the current sample of individuals with preserved linguistic abilities and low or without intellectual disability a modulation of STS activity related to face perception could be maintained or recovered. It should be noted, however, that the spatial distribution of the recorded activity was still atypical, with maximum activity observed in the middle of the face over the nose region. This result was surprising since this is a region deemed poor in social information and which, unlike the eyes, eyebrows or the mouth is not related to the perception of biological movements. An initial interpretation would be that the observed activity may represents an attempt at holistic face-processing, with the maximum amount of information being obtained by directing one’s gaze to the center of the faces. An additional interpretation would be that this pattern of activity is analogous to that produced by neurotypical subjects, but without the eye region.

**Figure 6.**
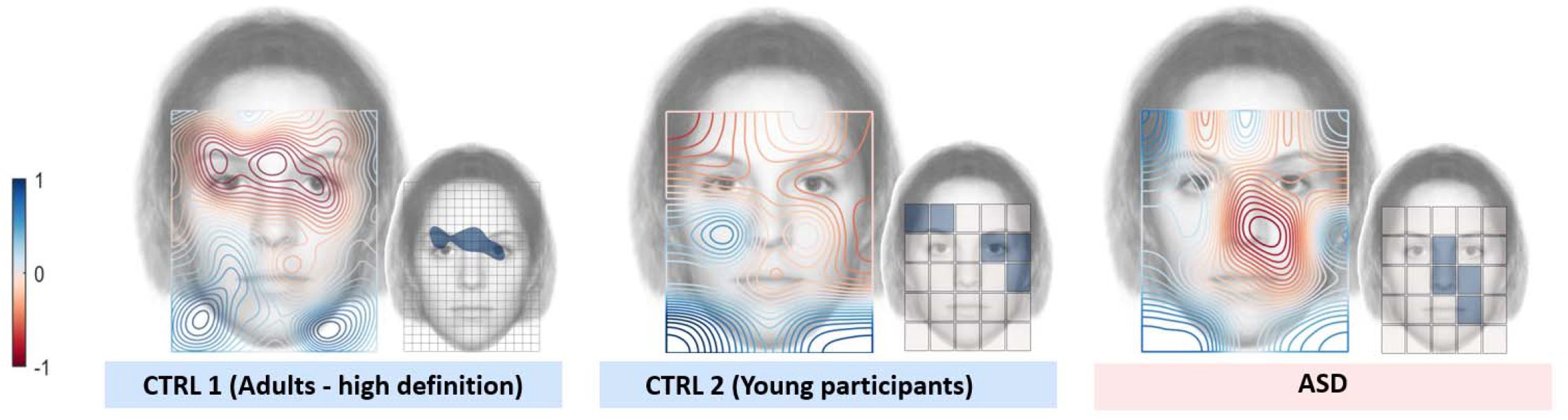
Evoked activity modulations in superior temporal regions are replicable in the control population, and atypical for ASD patients. Single trial modulations of the spatially filtered EEG, according to the face region focused on by subjects from respectively the Experiment 2 with a neurotypical population (left), Experiment 3 with a second population of young neurotypical participants (middle), Experiment 3 with ASD patients. On the large faces, the normalized spatial modulation of evoked activity in the STS at the group-level represented with a hot/cold contour plot. On the second face, regions where the expected activity is statistically significant (p<0.05, FWER corrected) are represented with a dark-blue color. Both neurotypical populations have produced the same response pattern with significant evoked activity in the area around the eye sand eyebrows that gradually decreases to reach two local minima on the lower left and right parts, where the focused area is outside the face. In the ASD population, at the group level, a significant modulation was also being observed but with a particular response pattern. The maximum and significant activity was no longer recorded in the eye area but rather at the center of the face, over the nose region.

## Discussion

Using EEG and spatial filtering techniques, we found that neurotypical subjects, but not participants with ASD, produced a unique evoked activity that was particular to the experience of making eye contact with natural face stimuli within a short interpersonal distance. This neural source was composed of a pronounced bilateral evoked activity in the superior temporal regions peaking maximum 240ms after stimulus onset. Moreover, we demonstrated that the particular region of the face stimulus that is presented to the fovea of subjects exerts a significant modulatory effect on this activity. Specifically, we observed maximal activity when the area of subject focus on was located between the two eyes of the face stimulus. For individuals with ASDs, who were exposed to the same viewing experience, we found this effect to be deficient. Interestingly the modulation of the STS signal we observed in the neurotypical population is reminiscent of the pattern of neural activity already reported in non-human primates. Mosher et al. (2014) have shown the existence of neurons that detect eye-contact in macaques monkeys’ amygdala. The maxima of these neurons’ activity was recorded during eye contact between the monkey’s observer and the monkey being observed in a video^59^. The face context designed in our study seems similar to the context that triggers maximal neuronal activity in the monkey’s brain and both suggest that rise and fall of neural activity is dependent on perception of eyes entering the subjects’ fovea field. Hence, our findings demonstrated that also in humans there may be dedicated cortical mechanisms that prioritize neural activity towards attention to eyes. This it may be a core disturbance of autism spectrum disorders.

Functional Magnetic Resonance Imaging (fMRI) studies have identified three cortical areas that appear to represent the core of the adult face processing system. These are the inferior occipital gyrus (‘OFA’ – occipital face area), the middle fusiform gyrus (FFA – Fusiform Face Area) and the superior temporal sulcus (STS)^35,58,60,61^. The current dominant neural models suggest that these structures are divided along two neural pathways of face processing. The ventral pathway, which includes the FFA, is more heavily implicated in the treatment of face traits. These are invariant facial aspects such as gender, age and identity. While the dorsal pathway, which includes the STS, is more specialized in processing the face state, this is made up of the changeable facial related processes such as speech, implied motion, emotional expressions, attention and intentions. This pathway is also considered as the gateway to an extended system dedicated to social perception encompassing the orbitofrontal cortex and the amygdala^53^61. Using a blind source separation method, we observed atypical evoked activity bilaterally in the STS region. In light of these results we can make a strong argument for the presence of an impairment of this second pathway in our ASD population.

STS has already been theorized as a keystone structure underlying the social-interaction impairments and verbal and non-verbal communication deficits which are characteristic of ASD^53,54,62^. STS abnormalities related to ASD have been observed systematically across numerous strata including anatomical^7–10^, metabolic^11–13,63^, structural^14^, and functional^64–66^. More specifically, previous work has demonstrated that in both hemispheres a region of the mid-posterior STS corresponding to the faces/voices regions (according to a standard anterior-posterior organization in the STS for different social tasks^67,68^), shows particular impairment in ASD patients^11–13,65^. This structure corresponds precisely with the cortical area detected in the current study. Through the application of appropriate signal processing techniques our work demonstrates that it is possible to observe, with high-density EEG, the dynamic of this strategic brain structure over a single trial. By demonstrating the electrophysiological eyes-sensitivity of the STS here, we have described a non-invasive approach that can be used to investigate a whole spectrum of possible electrophysiological modulations of the STS across a variety of tasks, clinical populations and neuropsychiatric conditions.

Moreover, we have not only demonstrated where an electrophysiological process critical for social perception, impaired in the ASD population, is located. We have also shown when this process occurs (≈240ms post-stimulus onset). The temporal location of this latency, following the well-known N170 component, does not come as such a surprise considering the estimated cortical source of the component. Indeed, intracranial recording studies using grids placed on the cortical surface, or intracerebral electrodes implemented within brain structures, have estimated the chronometry of activations evoked by the presentation of a face stimulus across the various structures of the ventral visual pathway. Recorded activities were mostly restricted to the fusiform gyrus prior to 200ms. Next, important stages of distributed parallel processes arose at 240 and 360 ms^25,36^, involving several regions located more anteriorly in the temporal cortex. Initial recognition specific processes also occur at these latencies (240ms), to up to 400-600ms within the hippocampus^36^. In this context, a latency occurring between 200 and 300ms is compatible with an evoked activity in the middle portion of the superior temporal sulcus.

The literature on how face processing develops (for a review see^61^) highlights a number of neural components of basic social perception (eventually assessed by the eyes-sensitivity of such component), that should be detected during the very early childhood. These include a preference in newborns for face-like stimuli over non-face stimuli^69–71^, a subsequent preferences in infants two months later to focus attention on the eyes of others^2,72^ as well as a tendency in early childhood to imitate the facial movements of others^73^ and to look at certain stimulus-general configurations that that resemble faces^74^. All of these behavioral observations suggest an early involvement of the superior temporal regions in the development of social cognition. Neuroimaging studies investigating the development of face processing suggests a similar hypothesis. While fMRI, PET and Near Infrared Spectroscopy techniques have revealed face-specific activations in superior-temporal regions for infants that are only months old^75,76^ fMRI studies have encountered difficulties in observing similar activations for children below ten-year-olds^77,78^. EEG studies throughout childhood present a similar pattern whereby only late detections (>250ms) of face specific components have been found before one year-old of age^79–83^ while an adult-like occipito-temporal N170 component can only be detected in older children (4-year-olds)^84^. Together, these results suggest the potential detection of an eye-sensitive electrophysiological component in superior temporal regions during early infancy, possibly even the first months of life. Such finding offers a critical new perspective on how we view the role of eye-contact behavior development^2^ and the involvement of the STS^54^ in the emergence of the spectrum of social disabilities known as “autism”.

## Materials and methods

### Experiment 1

#### Experimental Design

The first experiment consisted of random presentations of natural faces with neutral expressions. Subjects were instructed to focus on a fixation cross on the screen located at a point between the two eyes of each face stimulus. Each picture was presented for a duration of 167ms, with an SOA (Stimulus Onset Asynchrony) of 750 ±250ms (Fig 1). In order to ensure that subjects were paying attention throughout the task a blue dot was presented at random (p=0.0625), instead of a face stimulus. Subjects were instructed to press a button with their right index every time the blue dot appeared on the screen. We presented a total of 240 faces stimuli to at each participant.

#### Participants

We recruited a group of 13 adults (10 men and 3 women, mean age = 32, range = 22–50) with a clinical diagnosis of Asperger syndrome (AS) or high-functioning autism (HFA) according to Diagnostic and Statistical Manual-Revision 4 (DSM-IV R)^5^ and ASDI^85^. ASD participants were recruited from the nearby Vinatier psychiatric Hospital. Parents or caregivers were interviewed using the ADI-R^86^ to confirm the diagnosis (ADI-R Social : mean=16.5, σ=7.1 - ADI-R Communication-verbal : mean=8.87, σ=4.3 - ADI-R Repetitive behaviors : mean=3.5, σ=2.3). As part of this checking process, the French translation of A-TAC (autism, tics, AD-HD, and other comorbidities)^87^ was completed by the parents. Patients completed verbal and performance IQ tests (WAIS-III) and all showed average to above average estimates of intelligence (mean IQ=101, σ=16.2). We also tested a control group of 10 neurotypical subjects (5 men and 5 women, mean age = 26, range = 25–27) with normal or corrected to normal vision and without history of psychiatric or neurological disease. The protocol was approved by the Ethical Committee Sud-Est IV, Lyon.

#### Stimuli

We used a total of 12 faces as stimuli, in the study (3 men, 3 women and their horizontal-flip counterparts). These were built from built from photographic images selected from the Nimstim database^88^. We modified the proportions of the faces slightly using Gimp software (http://www.gimp.org/) so that we could control the distances between facial features for each picture. Specifically, the following factors were held constant: Interpupillary distance = 6.5°, eyes/nose distance = 5°, nose/mouth distance = 2°, mouth/chin distance = 5° - screen resolution ~31 pixels/degree of visual angle (Fig. S1). We deliberately choose pictures made up of real faces because of previously reported as critical stimuli for testing eye gaze abnormality in autism.^40^. Finally, interpersonal distance between subject and stimuli was set as relatively close in the context of standard personal space regulation (~56 cm - ^47,48^). Face stimuli angular size corresponds to this distance.

#### EEG recording and preprocessing

Continuous electroencephalogram (EEG) data was recorded from 64 AgCl carbon-fiber coated electrodes using an Electric Geodesic Sensor NetH (GSN300; Electrical Geodesic Inc., Oregon; http://www.egi.com/) with impedances kept below 50 kOhms. EEG data were recorded at a sampling rate of 1000 Hz with an online reference at the vertex electrode (Cz). One control subject (1 woman) was rejected during this step due to a technical-problem detected during the recording. Offline, data were band pass filtered using zero-phase Chebychev type II filters (Low pass - cutting frequency: 45 Hz, transition band width: 2 Hz, attenuation: 80 dB; order: 35, sections: 18 | High pass – cutting frequency: 0.3 Hz, transition band width: 0.2 Hz, attenuation: 80 dB; order: 9, sections: 5) and re-referenced to common average. Then, we epoched data from 200 ms before stimulus onset to 400 ms after stimulus onset. Corrupted epochs were automatically detected and rejected using EEGLAB^89^ and the FASTER toolbox^90^. Finally, for each subject, non-brain artifacts (eye movements, ballistocardiac noise, sensors movements and other electrical noises) were detected and rejected using independent component analysis (ICA) / blind source separation (BSS) with UW-SOBI (101 times delays). The UW-SOBI algorithm^91^ is an adaptation of the well-known SOBI algorithm^51,52,92^ reformulated as an uniformly-weighted nonlinear least squares problem to avoid the common “whitening” phase which is known to limit the performance of BSS/ICA algorithms in noisy conditions^93^.

#### Significance testing

We systematically used non-parametric statistical inference, which does not make assumptions about the distribution of the data, in the present study. The sample size was 22, 13 ASD participants, 9 controls subjects. All tests were two-sided and the familywise error rate was controlled with cluster-size inference when a multiple testing procedure was used (threshold at p<0.05 for cluster constitution, and p<0.05 for the cluster-corrected p-value)^94^.

#### Preliminary electrode space ERP analyses

We carried out two analyses which enabled to localize in time the neural evoked activities necessary to dissociate the ASD population from control subjects. First, we performed a simple topographic analysis which searched when and where differences in evoked power measured on the scalp could be assessed. Second, we used a multivariate pattern classification to estimate the classification power of the evoked pattern recovered by surface electrodes.

Thus, stimulus evoked activity was processed for each participant and at each electrode. Then, we assessed statistical differences between recovered evoked activities for the two populations using non-parametric Wilcoxon Rank-sum tests on each time-sample and each electrode, corrected by cluster-based permutations^94^. In a second step, we applied a multivariate pattern classification to determine when face processing in ASD patients could be reliably dissociated from face processing in control subjects. In this way, a classifier was trained and validated at each time-sample so that we could localize in-time when the best classification accuracy could be attained (Fig. 2A). To this aim, we used linear classification techniques^95,96^ using a 2-norm soft-margin support vector machine (SVM) with a linear kernel and a leave-one out cross validation procedure (MATLAB *‘svmtrain’* function with a soft margin parameter *‘boxconstraint’*=1). The classification accuracy is a noisy measurement and the procedure can lead, by chance, to overly high classification results. Therefore, the time-course of estimated accuracies (one by sample) were smoothed with a 40ms moving-average window. Moreover, to obtain a reliable estimate of the classification power of the evoked scalp patterns we used a bootstrap procedure. Thus, 100 time-courses of classification accuracy were estimated and averaged, using 150 randomly selected trials for each subject and each iteration for the estimation of the scalp patterns. Finally, we applied the minimum-noise forward model estimate^42^ in order to visualize the scalp patterns that lead to the best classification results (Fig. 2C).

#### Group Blind Source Separation

Group blind source separation (gBSS) offers a straightforward and computationally tractable solution to the problem of multi-subject analysis by creating aggregate data containing observations from all subjects. By providing a single estimation of the mixing and demixing matrices for the whole group, this strategy allows direct estimation of the components that are consistently expressed in the population. We employed UW-SOBI, a Second Order Statics (SOS) based BSS algorithm based on the approximate joint diagonalization of lagged-covariance matrices. This method is robust with respect to anatomo-functional inter-subjects’ variability^45^ and can separate group specific sources with non-proportional power-spectra without deleterious prior dimension reduction^44^. The other potential benefits of SOBI methods are first, their ability to separate correlated sources^51,52^ and second, a better sensitivity to the detection of critical sources that are often occulted by the most energetic phenomena^97^. One hundred one lagged-covariance matrices with time delays from 0/1000s to 100/1000s were calculated for each epoch. Then, lagged-covariance matrices were averaged across the dataset epochs first and then across subjects, resulting in 101 averaged lagged covariance matrices for the patient group and the control group. Finally, lagged covariances for both patients and control groups were averaged and approximately joint-diagonalized with the UWEDGE algorithm^98^, leading to the identification of 64 Independent Components (ICs) (Fig. S3 – 1a).

#### Sources spaces ERP analyses

In order to capture differences in how neurotypical subjects and ASD patients process natural neutral faces with neutral expressions, we processed stimulus evoked activity for each participant/ each electrode and each source. We tested statistical differences between evoked activities in the source space for each source and each time-sample using non-parametric Wilcoxon Rank-sum tests, corrected by cluster-based permutations^94^ (Fig. S3 – 1b).

#### Independent components sources localization

We used sLORETA software^99^ to estimate the intracerebral electrical sources separated by the BSS algorithm (Fig. S3 – 1c). The head model used for this analysis was obtained by applying the BEM method to the MNI152 template^100^. The 3D solution space was restricted to cortical gray matter and was partitioned into 6239 voxels with a spatial resolution of 5 mm. Then, the sLoreta solution to the inverse problem was computed using an amount of Tikhonov regularization optimized for an estimated Signal/Noise Ratio of 10 (Fig. S3 – 1d).

### Experiment 2

#### Experimental Design

We designed a second protocol with the aim of measuring the face region sensitivities of the source identified in the first experiment. This second task was developed specifically to assess single trial evoked response in the STS cortical region of interest identified in the first experiment. We recruited 14 neurotypical subjects (10 men and 4 women, mean age = 25, range = 19–31) with normal or corrected to normal vision and without history of psychiatric or neurological disease for this second experiment. A total of 2800 face stimuli were presented to each subject (2 sessions of 1000 presentations, one session of 800 presentations, stimulus duration: 167ms, SOA: 800 ± 100ms). Participants were instructed to focus on a fixation cross located on the center of the screen. Each stimulus consisted of a natural face with a neutral expression which was, displayed at a random location on the screen, multiplied by a Gaussian apodization window centered on the fixation cross (Fig 4). Thanks to this procedure, we were able to control the face region focused on by the subject as well as the luminance distribution projected onto their retina. After every 7 ± 3 stimuli were presented, a question mark appeared on the screen instead of a face stimulus. For these trials, participants had to recall the gender of the last displayed face and indicate their choice by pressing a button with their index or their middle finger. We included these trials as a mean of ensuring that participants maintained their attention throughout the task.

#### Stimuli

Face stimuli were identical to those used in the first experiment. However, for this version of the task, their location on the screen relative to the fixation cross was randomly determined for each trial. This method allowed each face region to be observed by the subject. With base coordinates [0°; 0°] located between the two pupils of each face picture, each observed region was drawn from a uniform distribution rectangle encompassing the whole face area (width [−8°; +8°], height [−12°; +8°] (Fig 3 - B). Considering the 2800 stimuli presented, the sampling density of face pictures was 2800/320=8.75 trials / degree^2^ / subject. To control the luminance distribution of visual stimuli, despite the different locations of face pictures on the screen, each stimulus was multiplied by a 2-dimensional Gaussian apodization function centered on the fixation cross. The width of the aperture window was set large enough (Full Widths at Half Maximum =10° of visual angle) so to ensure that the recognition of the gender or the identity of each stimulus was easy for all trials (Fig. 4) (Recognition rate = 98% - SD=2.9%).

#### EEG recording and preprocessing

We used Brain Product™ actiCHamp system to record the electroencephalographic signal from 128 active electrodes (actiCAP 128Ch Standard-2) mounted on an elastic cap at 10-10 and 10-5 system standard locations^101^. All electrode impedances were kept below 50 kOhms. Subjects were seated in a darkened, shielded room with their head position controlled by an ophthalmic chin-rest device so that their eyes remained at the same level of the fixation cross. EEG data were recorded at a sampling rate of 5000 Hz with an online reference at the Fz electrode. We band pass filtered data offline using zero-phase Chebychev type II filters (Low pass - cutting frequency: 45 Hz, transition band width: 2 Hz, attenuation: 80 dB; order: 35, sections: 18 | High pass – cutting frequency: 0.3 Hz, transition band width: 0.2 Hz, attenuation: 80 dB; order: 9, sections: 5) and re-referenced to a common average. Then, we epoched data from 200 ms until 400 ms after stimulus onset.

#### Single trial spatial filtering

In order to measure the single trial behavior of the EEG component identified in the first experiment, a spatial filter was calculated for each trial using minimum variance beamformer techniques^55,102^ in combination with the spatial information estimated at the group level with gBSS as described elsewhere^103^ (Fig. S3–2b). First, the mixing vector A_i_ of the IC of interest, estimated in the first experiment, was interpolated to the 128 electrodes layout using the scalp surface of the MNI152 template^100^ (Fig S3-1e).

Then, given the measured signal *x_t_* at trial *t*, the source signal 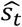 was estimated by:

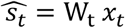

where the spatial filter *W_t_* is estimated by:

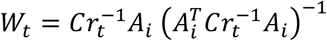

and the regularized noise covariance matrix *Cr_t_* by:

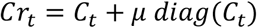

where *C_t_* is the data covariance matrix computed for the trial *t*, *μ* the Backus-Gilbert regularization parameter (=10) and *diag*(*C_t_*) the matrix of the diagonal elements of *C_t_* (the diagonal matrix of sensor noise).

#### Gaussian Kernel Density Mapping

We built, a source spatial sensitivity map for each subject using Gaussian kernel density mapping. Gaussian Kernel Density mapping is a non-parametric method for estimating the probability density functions of a random variable, weighted here by the intensity of the evoked activity estimated at the source level. The method is similar to ‘heat map’ representations used in eye-tracking studies (see e.g.^104^). At each trial *t*, where the area focused on located at the coordinates [x_t_, y_t_] on the face pictures, the mean evoked potential *m_t_* extracted from the cortical source of interest 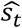, during the [200ms, 300ms] previously identified time-cluster, was multiplied by a two dimensional Gaussian kernel function with a mean value of [x_t_, y_t_] and a Full Width at Half Maximum (FWHM) of ~3.53° of visual angle (standard deviation = 1.5°). Then, the subject-level source spatial sensitivity map was built by averaging all Gaussian kernel functions. Finally, to highlight the ‘most positive’ and the ‘most negative’ areas, the mean value of the map was removed (Fig S3-2c).

For visualization purposes only, so that we could highlight the face areas that evoked the ‘most negative’ activities at the subject level, we thresholded each individual map using a non-parametric random-permutation test (p<0.05) (Fig 3 –A; Fig S3 – 2d).

Finally, we identified the face regions that the source was most sensitive to at the group level using the 14 maps estimated at the subject level. Each voxel was tested using the non-parametric, one tailed, sign test (306405 tests, p<0.05) while the Family Wise Error Rate (FWER) was controlled using the maxT/minP multiple testing procedure^105^, leading to a statistical non-parametric mapping of the evoked activity in the superior temporal regions (Fig 3 –B; Fig S3 – 2f). For all permutation-based tests the permuted values were, at the subject-level, the *m_t_* values.

#### Face regions of interest analysis

Since non-parametric testing on Gaussian Kernel Density maps can be overly conservative, we performed a second analysis based on the face regions of interest. We built three regions of interest (ROI): (1) a ‘Rectangular Eyes region’ (5° height, 12° width) encompassing the region around the eyes, (2) a ‘Triangular Nose/Mouth region’ (10° height, 12° width) encompassing the lower facial features and (3) a ‘no-facial feature region’ encompassing the other tested areas. For each subject, we calculated the Z-transformed mean evoked activities extracted from the cortical source of interest, during the [200ms, 300ms] time-cluster at each ROI. Then, we processed a group analysis using the Kruskal-Wallis non parametric test with post-hoc multiple comparisons FWER corrected by the Tukey-Kramer method.

### Experiment 3

#### Experimental Design

In Experiment 2b we replicated Experiment 2a by introducing minor modifications to adapt the protocol for two populations. The first population was composed of patients who had been diagnosed with Autism Spectrum Disorders while the second population comprised of young neurotypical controls. Thus, for this second experiment we recruited 14 neurotypical subjects (5 males and 9 females, mean age = 11.6, range = 6–21) with normal or corrected to normal vision and without history of psychiatric or neurological disease and 14 ASD patients (14 males, mean age = 20, range = 18–21) with normal or corrected to normal vision.

Patients from the ASD group were recruited from the Azienda Ospedaleria Brotzu (Cagliari, Italy) and had a clinical diagnosis of autism according to the *Diagnostic and Statistical Manual of Mental Disorders, 5^th^ edition*^5^. At the time of the study all patients had been rehabilitated and exhibited a broad range of intelligence estimates (WAIS-IV^106^: range [58-141], mean: 97) and autistic symptoms intensity (ADOS^86^ – Mod IV : range [2-14], mean: 8). All participants had normal or corrected to normal vision. The study was approved by French (Sud-Ouest, project N° 2018-A02037-48) and Italian Ethical Committees (Azienda Ospedaliero-Universitaria of Cagliari, project N°AOB/2013/1, EudraCT code 2013-003067-59) and a written informed consent was obtained from all participants and/or their legal representative prior to their inclusion in the study.

The first modification made to the original study was to significantly reduce the number of trials presented to subjects in order to reduce the total duration of the experiment. Thus, a total of 1600 face stimuli were presented to each subject (1 training session of 100 presentations – unrecorded – then 3 sessions of 500 presentations, stimulus duration: 167ms, SOA: 800 ± 100ms).

Second, unlike the previous version, the region focused on for each stimulus presentation was no longer randomly selected from all the pixels that made up the face region but instead was randomly selected with a uniform distribution from 25 predetermined locations or regions of interest (ROI). We made this adjustment in order to optimize the statistical power of the analysis given the reduced number of trials. ROIs were constituted by dividing the region of the studied face stimuli (20° height, 16°width) by a grid of 5×5, each ROI being a rectangle of size height=4° x width=3.2°.

Given the 1500 trials presented and recorded, the sampling density for face pictures was 1500/25= 60 trials / ROI / subject. Finally, we obtained the spatial sensitivity map of the investigated electrophysiological source by performing all signal and statistical analyses on the 25 locations (25 non-parametric, one tailed, sign tests, p<0.05, FWER controlled using the maxT/minP multiple testing procedure^105^).

For visualization purposes only, we obtained a smoothed representation of the results using bicubic interpolation. This allowed us to map variations in intensity of the evoked activity of interest as a function of the region focused on by subjects (Fig 6).

## Acknowledgments

This work was supported by Centre National de la Recherche Scientifique and by Labex Cortex University of Lyon I (“Investissement d’Avenir”) Grant ANR-11-LABEX-0042 to AS.

## Author contribution

AS and GL proposed the concept study and designed the research; GD and RF recruited patients and performed clinical evaluation. MC, GL and AS performed the EEG experiments. G.L. analyzed data. G.L. and A.S. wrote the paper. MC, GD, RF and CD provided helpful discussion during data analysis. All authors discussed data and approved the final manuscript.

## Supplementary Materials

**Figure S1.**
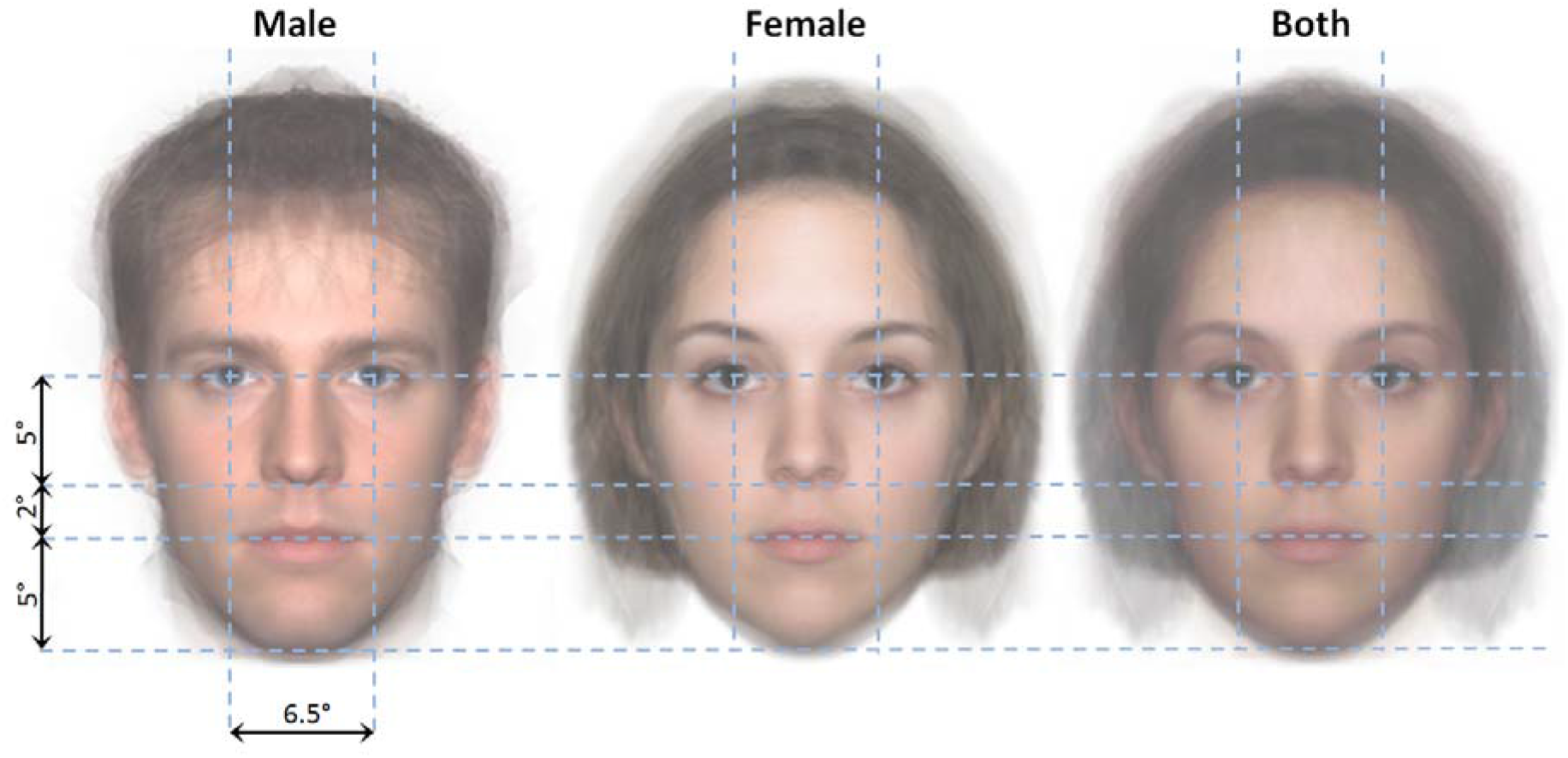
Stimuli. Average of respectively male/female/all stimuli used in this study. Proportions of the faces were controlled in order to maintain exactly the same distances between facial features for each picture: Interpupillary distance = 6.5°, eyes/nose distance = 5°, nose/mouth distance = 2°, mouth/chin distance = 5°. Real face pictures were chosen since previously studies have reported increasing sensitivity of ecological stimuli to detect gaze abnormalities studies in autism^40^. Finally, angular size of the face stimuli corresponded to a relatively close interpersonal distance in the context of personal space regulation (~56 cm - ^47,48^).

**Figure S2.**
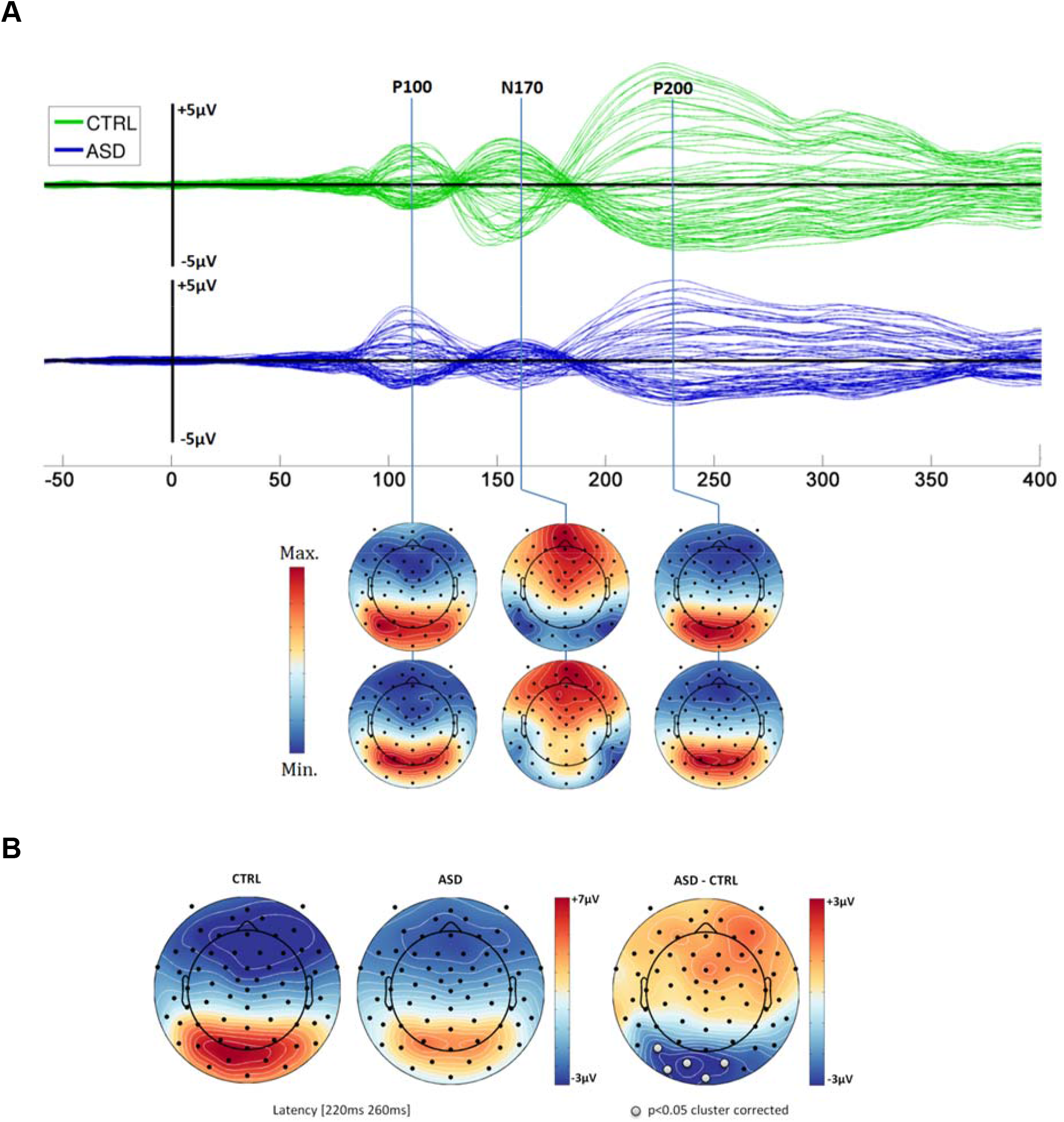
Surface ERP analysis. *Experiment 1:* Surface Event Related Potential (ERP) analysis. **A-** Butterfly plot and scalp topography of the face evoked activity for control (CTRL, n=9) and Autism Spectrum Disorder (ASD, n=13) population. Both evoked the classical P100, N170 ERPs. **B-** As predicted and discussed in recent studies and meta-analyses (e.g. ^33,34^), our results did not report a difference between ASD and control population for the amplitude of classical P100 and N170 ERPs (Wilcoxon rank-sum test p>0.05 uncorrected – subjects were instructed to focus on a fixation cross located between the two eyes of face stimuli). The only significant difference revealed in the electrode space in our study was located on a cluster of occipital electrodes, 220ms to 260ms after the stimulus onset (CTRL > ASD, cluster permutation test, p<0.05).

**Figure S3.**
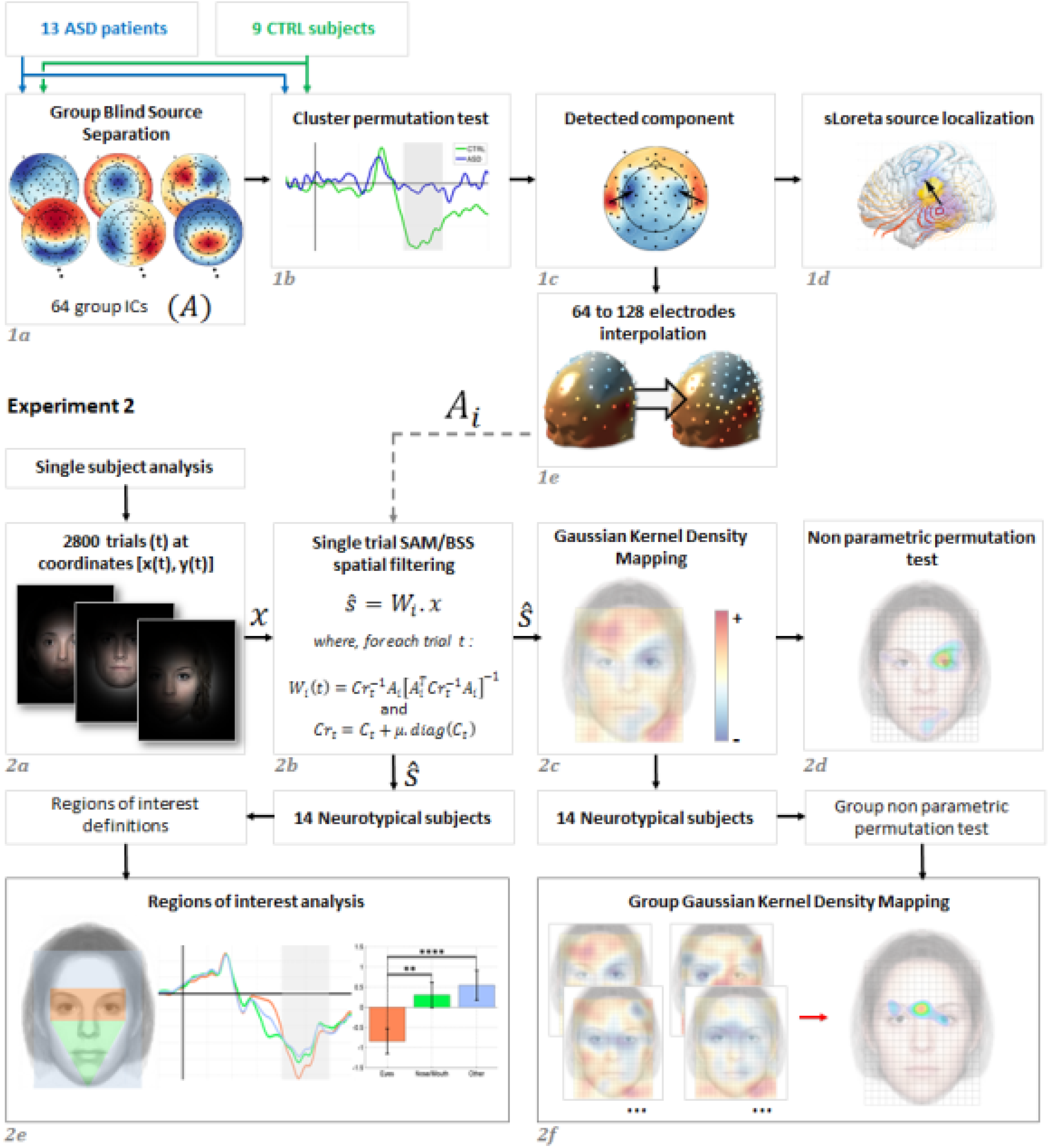
Signal processing pipeline. Signal processing pipeline for both experiments: **1a**– After Group Blind Source Separation (gBSS/gICA), 64 task specific components were estimated for the whole population. **1b**-Cluster permutation test revealed significant differences between the evoked activity of ASD patients and CTRL subjects at the source (Independent Components - ICs) level. **1c**– Scalp topography of the identified significant component was extracted from the weights of the estimated BSS demixing matrix. **1d**– Cortical localizations of the detected ICs were estimated with the sLoreta method^99^. **1e**– In order to transpose the spatial signature of the source of interest, for the 128 electrodes montage of the second experiment EEG device, the weights of the mixing matrix were spatially interpolated according to the head surface of the MNI152 template^100^. **2a**– For each subject, 2800 face stimuli were presented. For each trial t, the coordinates [x(t), y(t)] represented the location on the face stimulus of the area focused on. **2b**– Each trial was spatially filtered using the Synthetic Aperture Magnetometry technique (SAM^55^) and the BSS mixing vector estimated in the experiment 1^103^. **2c**– For each subject, a source spatial sensitivity map was built using Gaussian kernel density mapping. At each trial *t*, the mean evoked potential *m_t_* extracted from the cortical source of interest, during the [200ms, 300ms] time period, was multiplied by a two dimensional Gaussian kernel function with a mean value of [x(t), y(t)] and a Full Width at Half Maximum (FWHM) of ~3.53° of visual angle (standard deviation = 1.5°). Then, the subject-level source spatial sensitivity map was built by averaging all Gaussian kernel functions. Finally, to highlight the ‘most positive’ and the ‘most negative’ areas, the mean value of the map was removed. **2d**– For visualization only, the subject level threshold delimiting the most ‘source sensitive’ face regions was estimated by non-parametric permutation test (random perturbations of *m_t_* values, p<0.05). **2e -** Face regions of interest analysis. For each subject, the mean evoked activities extracted from the cortical source of interest, during the [200ms, 300ms] time period, were calculated at each ROI and Z-transformed. Then, a group analysis was processed using the Kruskal-Wallis non parametric test with post-hoc multiple comparisons FWER corrected by the Tukey-Kramer method. **2f**– Group analysis of the 14 subject’s source spatial sensitivity maps. Each voxel was tested using the non-parametric, one tailed, sign test (306405 tests, p<0.05) while the Family Wise Error Rate (FWER) was controlled using the maxT/minP multiple testing procedure^105^. The region around the eyes region presented the most powerful evoked activity.

**Figure S4.**
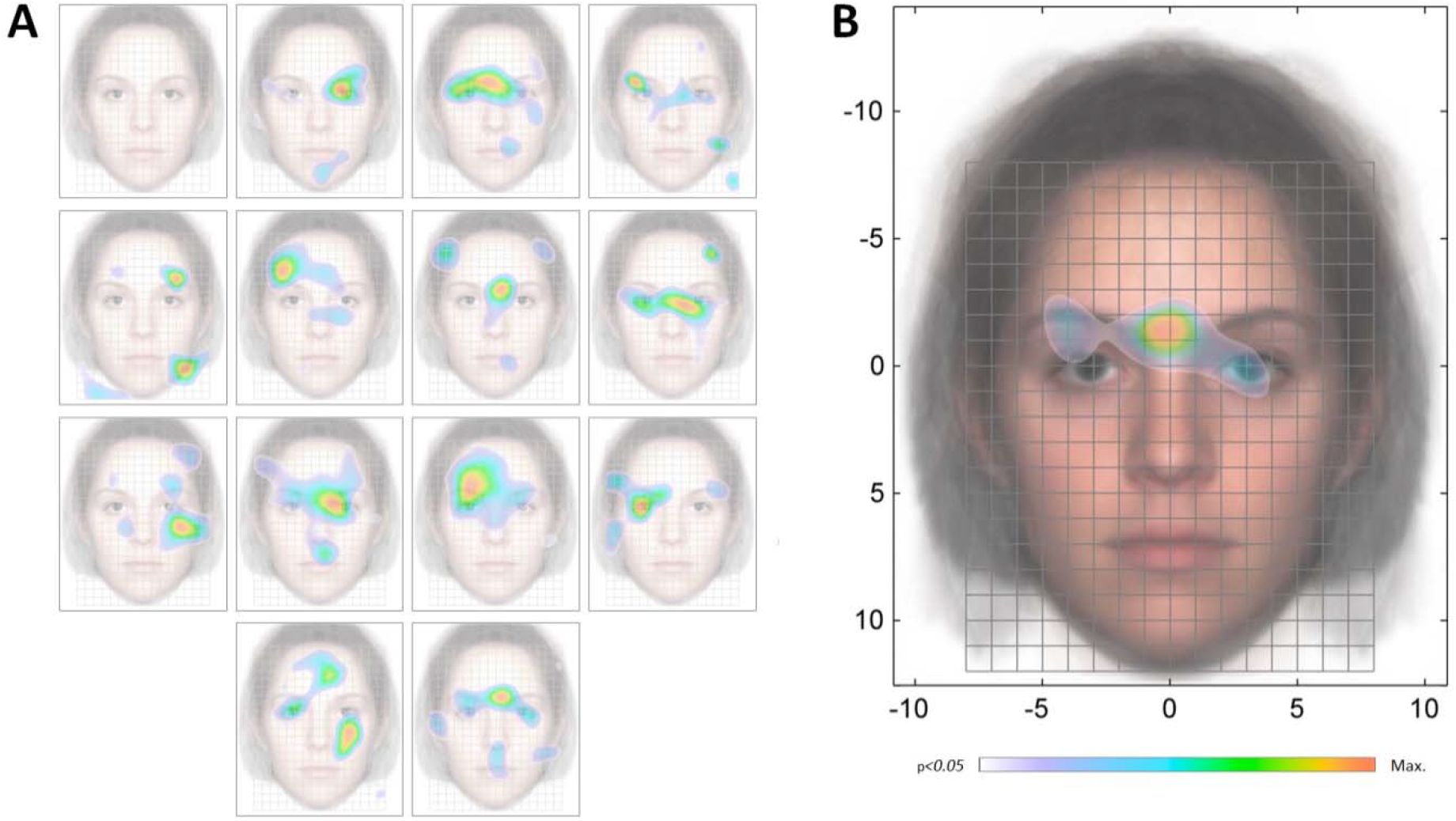
Statistical non-parametric mapping of the evoked activity in STS/STG. Gaussian Kernel Density mapping of the evoked activity of the spatially filtered EEG (STS/STG, [200ms 300ms] post-stimulus onset). **A**- Single-subject analysis (permutation test p<0.05). **B**– Group-level analysis (permutation test, p<0.05 FWER corrected).

